# *in silico* Assessment of Antibody Drug Resistance to Bamlanivimab of SARS-CoV-2 Variant B.1.617

**DOI:** 10.1101/2021.05.12.443826

**Authors:** Leili Zhang, Tien Huynh, Binquan Luan

## Abstract

The highly infectious SARS-CoV-2 variant B.1.617 with double mutations E484Q and L452R in the receptor binding domain (RBD) of SARS-CoV-2’s spike protein is worrisome. Demonstrated in crystal structures, the residues 452 and 484 in RBD are not in direct contact with interfacial residues in the angiotensin converting enzyme 2 (ACE2). This suggests that albeit there are some possibly nonlocal effects, the E484Q and L452R mutations might not significantly affect RBD’s binding with ACE2, which is an important step for viral entry into host cells. Thus, without the known molecular mechanism, these two successful mutations (from the point of view of SARS-CoV-2) can be hypothesized to evade human antibodies. Using *in silico* all-atom molecular dynamics (MD) simulation as well as deep learning (DL) approaches, here we show that these two mutations significantly reduce the binding affinity between RBD and the antibody LY-CoV555 (also named as Bamlanivimab) that was proven to be efficacious for neutralizing the wide-type SARS-CoV-2. With the revealed molecular mechanism on how L452R and E484K evade LY-CoV555, we expect that more specific therapeutic antibodies can be accordingly designed and/or a precision mixing of antibodies can be achieved in a cocktail treatment for patients infected with the variant B.1.617.

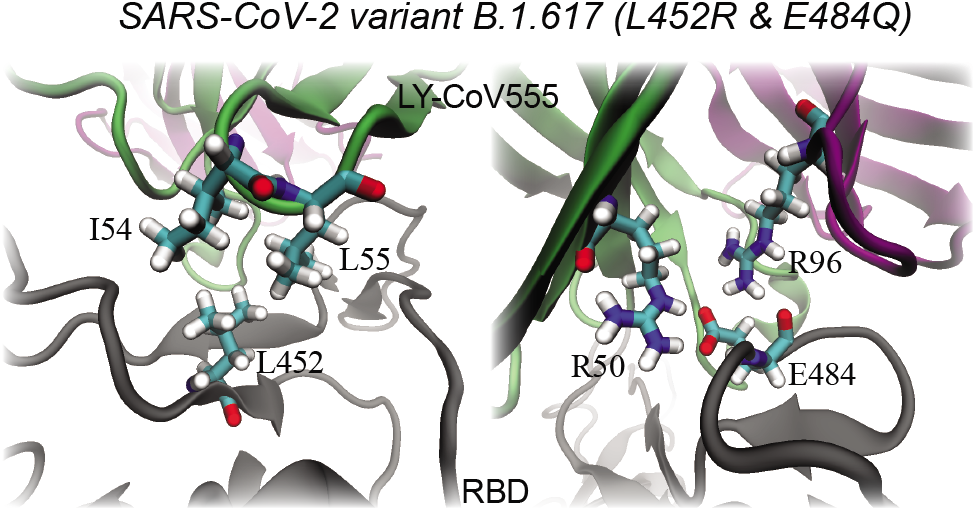

## Introduction

The severe acute respiratory syndrome coronavirus 2 (SARS-CoV-2) that causes the ongoing COVID-19 pandemic has evolved into several new dominant variants with major genomic changes through mutations. Mutations in viruses arise (partly) as a result of low polymerase fidelity of viral replication but become a survival mechanism for viruses to adapt to new hosts and environments.^1^ Although the majority of viral mutations are benign with most of them weeded out immediately, the current pandemic provides a suitable environment for SARS-CoV-2 to make natural selection of rare-acted but favorable mutations to strengthen its survival capability. Since the virus surface spike protein plays an important role in mediating SARS-CoV-2 entry into human cells and is the target for vaccine and therapeutic development, any mutations on this region may have biological significance as it could affect the viral infectivity and antigenicity.^2–4^ Indeed, experimental studies showed that the D614G mutation discovered at the earlier stage of the pandemic enhances the virus fitness and increases its transmission.^5,6^ Similarly, the N501Y mutation found in the B.1.1.7, B.1.351 and B.1.1.28.1 variants has been demonstrated to increase the binding affinity between the receptor-binding domain (RBD) and its human receptor ACE2 (hACE2), making these variants more transmissible.^7–9^ Moreover, experimental and computational studies have showed that the K417N and E484K found in the B.1.351 variant could evade neutralization by many monoclonal antibodies.^10,11^

Recently, a variant named B.1.617 that carries two mutations including the L452R and E484Q has been designated as variant of interest (VOI) by the World Health Organization (WHO), suggesting that B.1.617 potentially could have higher transmissibility, severity or reinfection risk and is required continuous monitoring. In fact, the two mutations found in B.1.617 are not completely new and have been seen in other variants separately. For example, the L452R mutation has been spotted in the B.1.427/B.1.429 variant which is known to be more contagious and is capable of escaping antibody neutralization.^12^ Also, the E484Q mutation is similar to the E484K found in the B.1.351 and B.1.1.28.1. The latter were found to reduce neutralization by convalescent antisera and binding of some monoclonal antibodies and increases the binding affinity to the hACE2.^10,11^ For the B.1.617 variant, this is the first time that these two mutations are found to coexist together, therefore it is important to understand how this variant could evade human antibodies or existing antibody drugs for treating COVID-19.

Complementary to ongoing experimental efforts, the all-atom molecular dynamics (MD) simulations with well-calibrated force fields have been widely used to explore the molecular mechanism of proteins.^13–16^ In this work, we carried out an *in silico* exploration on how a monoclonal antibody that can neutralize the wide-type SARS-CoV-2 but failed to target the B.1.617 variant. Here, we focus on the biologics drug LY-CoV555 that is a monoclonal antibody isolated from a convalescent COVID-19 patient. LY-CoV555 recognizes an epitope site in the RBD overlapping the binding site of ACE2, and was found to be efficacious on the wide-type SARS-CoV-2.^17^ By combining all-atom molecular dynamics, alchemical calculations and deep learning (DL) analysis, we aim to unveil the molecular mechanism on how these two mutations in the variant B.1.617 can be evasive to LY-CoV555. Our results provide invaluable insights for future designs of more efficacious mAbs to treat COVID-19 patients infected with the variant B.1.617, and highlight the need for precision medicine (biologics) for each SARS-CoV-2 variant.

## Results

### MD simulations of the complex of RBD and LY-CoV555

Figure 1a illustrates the simulation system for modeling the interaction between LY-CoV555 and RBD (see the Methods section for detailed simulation protocols). According to our previous simulation of a different complex of RBD and the Fab of the human antibody CB6,^9,18^ the constant domains C*^H^* in the heavy chain and C*^L^* in the light chain (connected through disordered coils with their respective variable domains V*^H^* and V*^L^*) are not relevant for studying CB6’s binding with RBD. Thus, in this work, we only include the V*^H^* and V*^L^* domains of the Fab of LY-CoV555 in our simulation system, as shown in Fig. 1a. This complex is very similar to the one in the crystal structure (PDB code: 7K43) showing that RBD is bound with a Fab fragment (V*^H^* and V*^L^*) of the antibody S2M11.^19^ Our modeled complex was solvated in a 0.15 M KCl electrolyte. Hereafter, we simply refer the variable domains that interact directly with RBD as LY-CoV555. Additionally, we refer the RBD of the variant B.1.617 as RBD-v.

**Figure 1:**
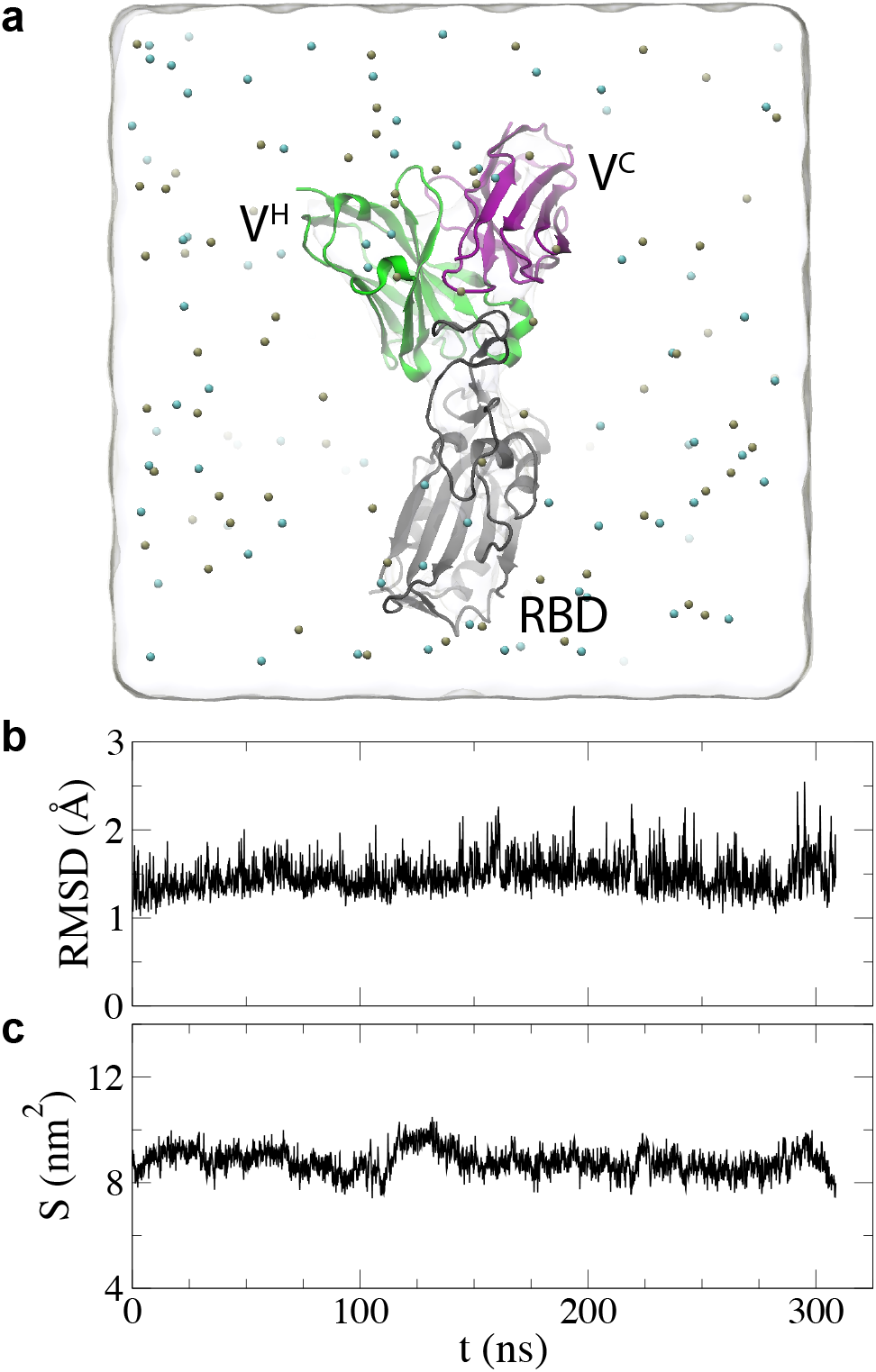
MD simulation systems. a) Simulation setup for the complex of wide-type RBD and LY-CoV555 (a Fab fragment). Proteins are in the cartoon representation, with RBD in gray, the V*^H^* domain in the heavy chain of LY-CoV555 and the V*^L^* domain in the light chain of LY-CoV555 in green and purple respectively. K^+^ and Cl^−^ are shown as tan and cyan balls respectively. Water is shown transparently. b) Time-dependent RMSD values for the complex (including only backbone atoms). c) Time-dependent contact areas *S* between RBD and V*^H^*/V*^L^* domains.

During the 300 ns or so MD simulation, the complex of RBD and LY-CoV555 starting from the structure in the crystal environment (PDB code: 7KMG) was properly equilibrated in the physiology-like environment (a 0.15 M electrolyte). Figure 1b shows the root-meansquare-deviation (RMSD) of all protein-backbone atoms in the complex. After about 50 ns, the RMSD values saturated at around 1.5 Å for the entire complex, suggesting that not only the secondary structure of each monomer (RBD, LY-CoV555’s V*^H^* or V*^L^*) but also the whole trimer-structure were very stable.

We further calculated the interfacial contact areas for the complex using the solvent accessible surface area (SASA) method. ^20^ By definition, the contact area is calculated by (SASA_LY–CoV555_+SASA_RBD_-SASA_complex_)/2. On average, Figure 1c shows that the contact area is about 8.8 nm^2^. During the entire simulation time, values of contact areas fluctuated around the mean value, corroborating that the complex structure was stable. These results highlight that LY-CoV555 is an effective neutralizing antibody for the wide-type SARS-CoV-2.

To evaluate whether LY-CoV555 is still effective for the variant B.1.617 carrying the L452R and E484Q mutations, we further investigate interfacial coordinations around L452 and E484. Remarkably, both L452 and E484 play an important role in stabilizing the interfacial binding, as shown in Figure 2. Figure 2a highlights the hydrophobic interactions between L452 in RBD and I54/L55 in V*^H^* of LY-CoV555, while Figure 2b signifies two interfacial salt-bridges: 1) between E484 in RBD and R50 in V*^H^* of LY-CoV555; 2) between E484 in RBD and R96 in V*^L^* of LY-CoV555. These favorable interfacial interactions provide molecular mechanism on how LY-CoV555 can be stably bound on RBD.

**Figure 2:**
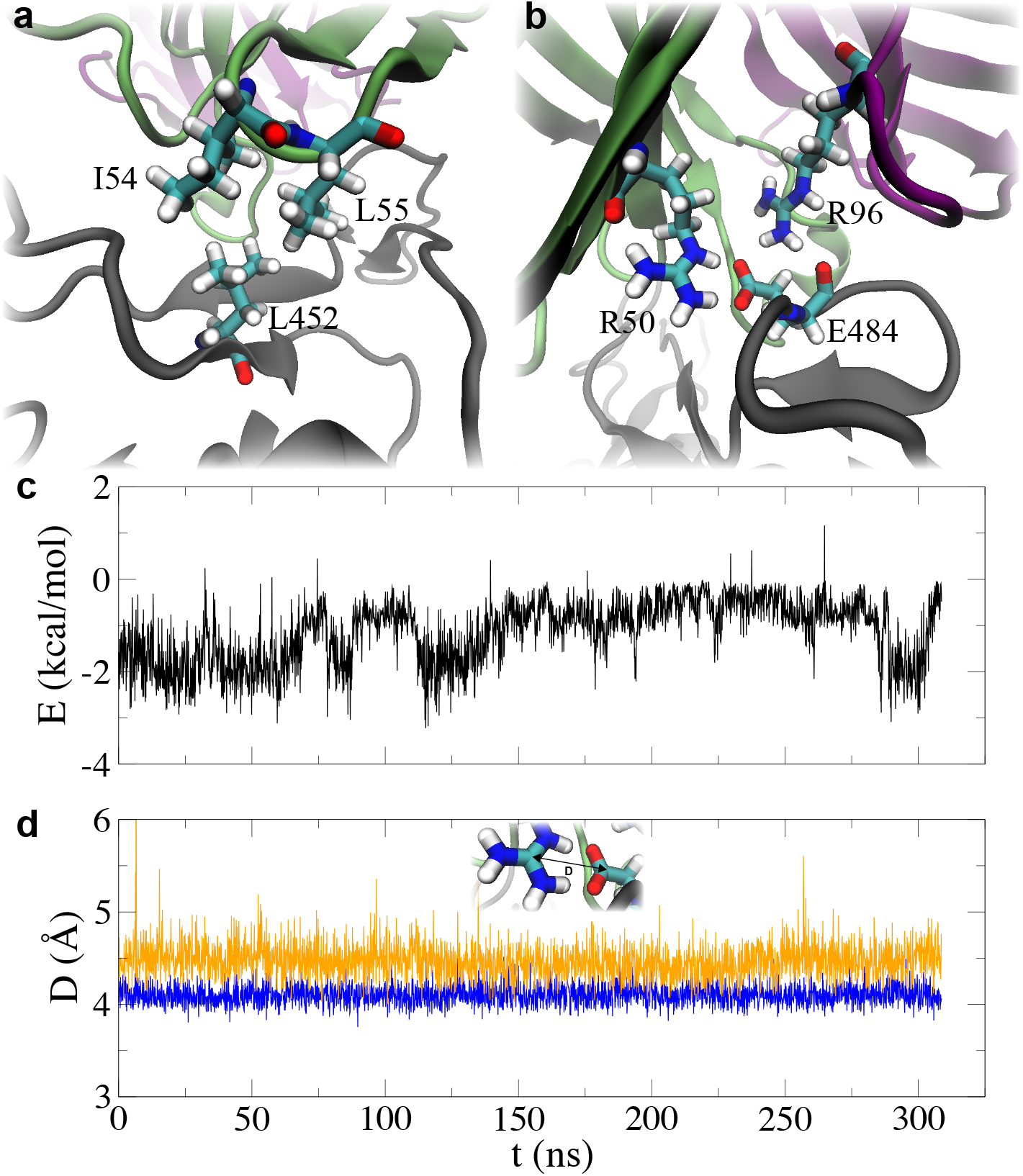
Favorable interactions between LY-CoV555 and RBD. a,b) Illustrations of favorable coordinations for L452 and E484 that are mutated into R452 and Q484 in the variant B.1.617. c) Time-dependent interaction energies between L452 in RBD and I54/L55 in the V*^H^* domain of (LY-CoV555). d) Time-dependent distances between the CZ atom in (R50 or R96) and the CD atom in E484. The inset illustrates the definition of the distance *D*. Results for the salt-bridges R50-E484 and R96-E484 are colored in blue and orange, respectively.

From the analysis of simulation trajectory, we found that the interaction energy between L452 and I54/L55 is dominated by the van der Waals interactions, which is understandable because of the hydrophobic nature. Figure 2c shows that during the simulation this interaction reduce the potential energy by 1.2 kcal/mol averagely. In addition, two energy levels can be discerned in the time-dependent interactions (Figure 2c), resulting from the fact that I54 can temporarily move away from L452 from time to time due to thermal fluctuations. When both I54 and L55 interacted with L452, the mean interaction energy is about −2 kcal/mol. However, with only L55 bound with L452, the interaction was weaker and the mean energy is only about −0.5 kcal/mol (Figure 2c).

To demonstrate that two salt-bridges shown in Figure 2b were stable, we calculated time-dependent characteristic distance *D*, defined as the distance between the CZ atom in R50 (or R96) and the CD atom in E484 (see the inset in Figure 2d). Figure 2d shows that the time-dependent distances *D* between R50 and E484 is nearly constant and the average distance is 4.1 Å. The salt-bridge between R96 and E484 has a different pose (Figure 2b), leading to a larger average distance of 4.5 Å. Noticeably, the fluctuation in time-dependent distances is larger for the salt-bridge between R96 and E484 than for salt-bridge between R50 and E484, suggesting that the former salt-bridge is relatively weaker. Overall, the local structure formed by E484, R50 and R96 was stable (Movie S1 in Supporting Information, because these two salt-bridges were actually buried inside the protein complex (Figure S1 in Supporting Information). Note that generally a water-exposed salt-bridge can form and break frequently and thus their distances can have two different mean values corresponding to formed and broken states of the salt-bridge.^9^

### Free energy perturbation calculations for L452R and E484Q mutations

Unfortunately, the importance of L452 and E484 in stabilizing the LY-CoV555’s binding with RBD as shown above also indicates that L452R and E484Q mutations can be highly evasive ones for LY-CoV555. Here, we performed free energy perturbation (FEP) calculations^21^ to obtain the binding free energy change (ΔΔ*G*) induced by the L452R and E484Q mutations. As required in FEP calculations, we performed 108-ns-long MD simulations of RBD in a 0.15 M KCl electrolyte (a free state), as shown in Figure S2a in Supporting Information. Interestingly, the saturated RMSD values for RBD’s backbone atoms are around 2 Å (Figure S2b in Supporting Information) that is larger than the one (1.5 Å) for the complex, which is due to the flexible coil (containing E484) in the free state (Movie S2 in Supporting Information). While in the bound state, the same coil is docked at the LY-CoV555’s interface (see Figure 2b) and thus less flexible. It is worth noting that due to the flexibility of the above mentioned coil, in the variant B.1.351 the mutated K484 residue can move toward E75 in ACE2 and form a salt-bridge,^9^ enhancing the binding affinity. With protein structures for both bound and free states in respective MD simulations, we applied the FEP alchemy method to obtain the binding free energy difference for L452R and E484Q mutations on the RBD. The protocol is briefly described in the Methods section and the detailed one can be found in previous works. ^9^

During a typical FEP calculation, the original and new residues gradually disappears and appears respectively. After the L452R mutation, Figure 3a shows R452 is in contact with hydrophobic I54 and L55 residues (low dielectric media) in LY-CoV555. However, in the free state, R452 is surrounded by water/ions (high dielectric media), which suggests that it is unfavorable for R452 to be at the interface between RBD and LY-CoV555. The rigorous FEP calculation yielded ΔΔ*G* of 3.04 kcal/mol, corroborating that the L452R mutation is energetically unfavorable. Our FEP result together with the role of L452 in the interfacial hydrophobic interaction (Figure 2a) provides theoretical explanations on why mutations of L452 to charged E, D, R and K residues are highly evasive for LY-CoV555 in experiment. ^22^

**Figure 3:**
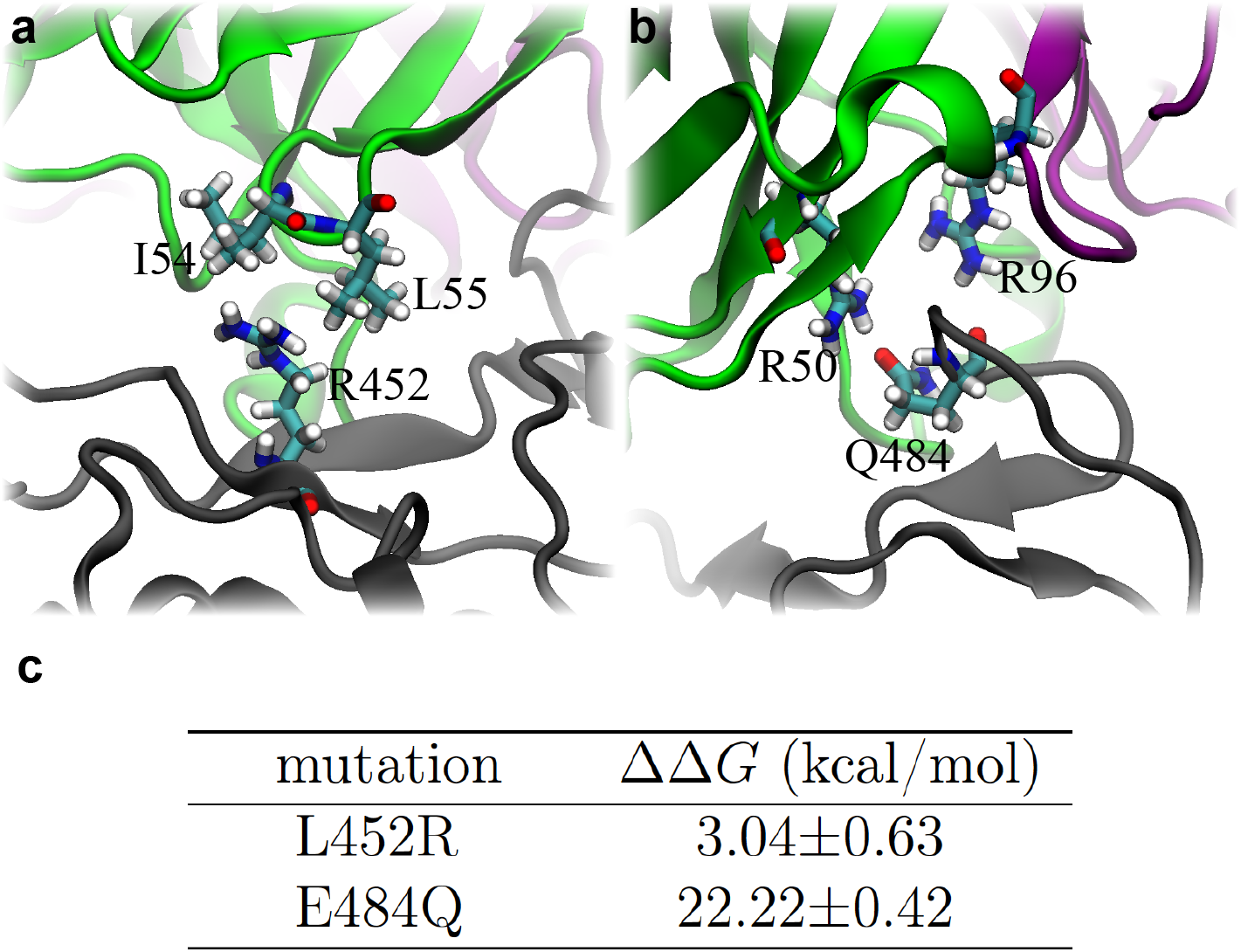
Interactions between LY-CoV555 and the variant B.1.617. a) Illustrations of unfavorable interfacial coordinations for L452R. b) Illustrations of unfavorable interfacial coordinations for E484Q. The RBD, V*^L^* and V*^H^* are colored in gray, green and purple respectively. Key residues at the interface are in the stick representation. c) Free energy changes for mutations L452R and E484Q.

Figure 3b shows that after the E484Q mutation Q484 was away from R96 and formed a hydrogen bond with R50. Compared with the local coordinations before the E484Q mutation (Figure 2b), the local interfacial interaction was significantly weakened after the removal of two salt-bridges that were buried inside the complex and stabilized the entire complex (See Figure S2 in Supporting Information). Overall, the E484Q mutation yields an extra charge (+e, where e is the elementary charge) buried inside the low dielectric protein-media, which is highly unfavorable from the free energy point of view. From the FEP calculation, ΔΔ*G* ~ 22.22 kcal/mol, indicating a much reduced binding affinity between RBD-v and LY-CoV555. Note that the result of ΔΔ*G* is seemingly large but is consistent with previous result that the removal of one buried salt-bridge yields ΔΔ*G* of about 10 kcal/mol.^9^ Taking all together, it is concluded that LY-CoV555 cannot bind RBD-v (of the variant B.1.617).

With the revealed molecular mechanism for RBD’s binding with LY-CoV555 and energetics for mutations, it becomes possible to design or engineer a more efficacious antibody drug targeting the variant B.1.617 specifically. Meanwhile, it is advantageous to stay one-step ahead of SARA-CoV-2 by considering what other possible mutations can be. Previously, we conducted an extensive *in silico* alanine-scan of all interfacial residues of RBD to determine those key residues that form favorable binding with the antibody CB6.^18^ Thus, mutations of those residues could significantly reduce the binding affinity between RBD and an antibody. In this work, we employed an efficient DL method (see below) to identify all residues in RBD that are important for the binding with LY-CoV555.

### CASTELO identifies L452 and E484 among the most beneficial residues in RBD while interacting with LY-CoV555

In previous works,^23,24^ we demonstrated that our designed MD-ML workflow CASTELO was able to identify atoms or atom groups (including amino acid residues) that contributed positively and negatively to the two-body interactions (such as drug-protein interactions^23^ and peptide-protein interactions^24^), agreeing with both experimental measurements and computationally expensive free energy calculations. Here, we applied the same workflow to the MD simulation for the complex of RBD and LY-CoV555. With the simulation trajectory, we first used RMSD clustering method to determine whether the simulation contained a stable binding structure between RBD and LY-CoV555. As shown in Figure 4A, with RMSD clustering method we identified that the largest cluster of the trajectory lasted ~169 ns far longer than the threshold of structural stability used previously (50 ns),^23,24^ which suggests that the simulation used in this study contains a stable binding structure of RBD and LY-CoV555. It is worth mentioning that this is an important first step. As we showed previously, the identification of beneficial or malicious residues relies on the comparison between the clustering results of the overall structures and the clustering of the individual residues. If the overall structures (the interfacial residues in this case) are stable, low similarity between the clusters of the stable overall trajectory and the clusters of an individual residue would mean that the individual residue is unstable. Residues such as these would be identified as ones that harm the interactions (which we call the malicious residues). On the contrary, high similarity between the clusters of the stable overall trajectory and the clusters of an individual residue would mean that the individual residue is stable. Residues such as these would be identified as ones that benefit the interactions (which we call the beneficial residues). However, if the overall structures are unstable, two problems would occur. First, in the simulations where no stable binding structures are verified, the confidence of finding the actual binding state is generally low. Therefore, predictions based on these simulations might be less meaningful. Second, comparison metrics (i.e. *dv* scores) may not differentiate between unstable residues and stable residues when the overall structure is unstable. In summary, verifying that our simulation contains a stable binding structure of RBD-LY-CoV555 is the prerequisite of the subsequent CASTELO analysis.

**Figure 4:**
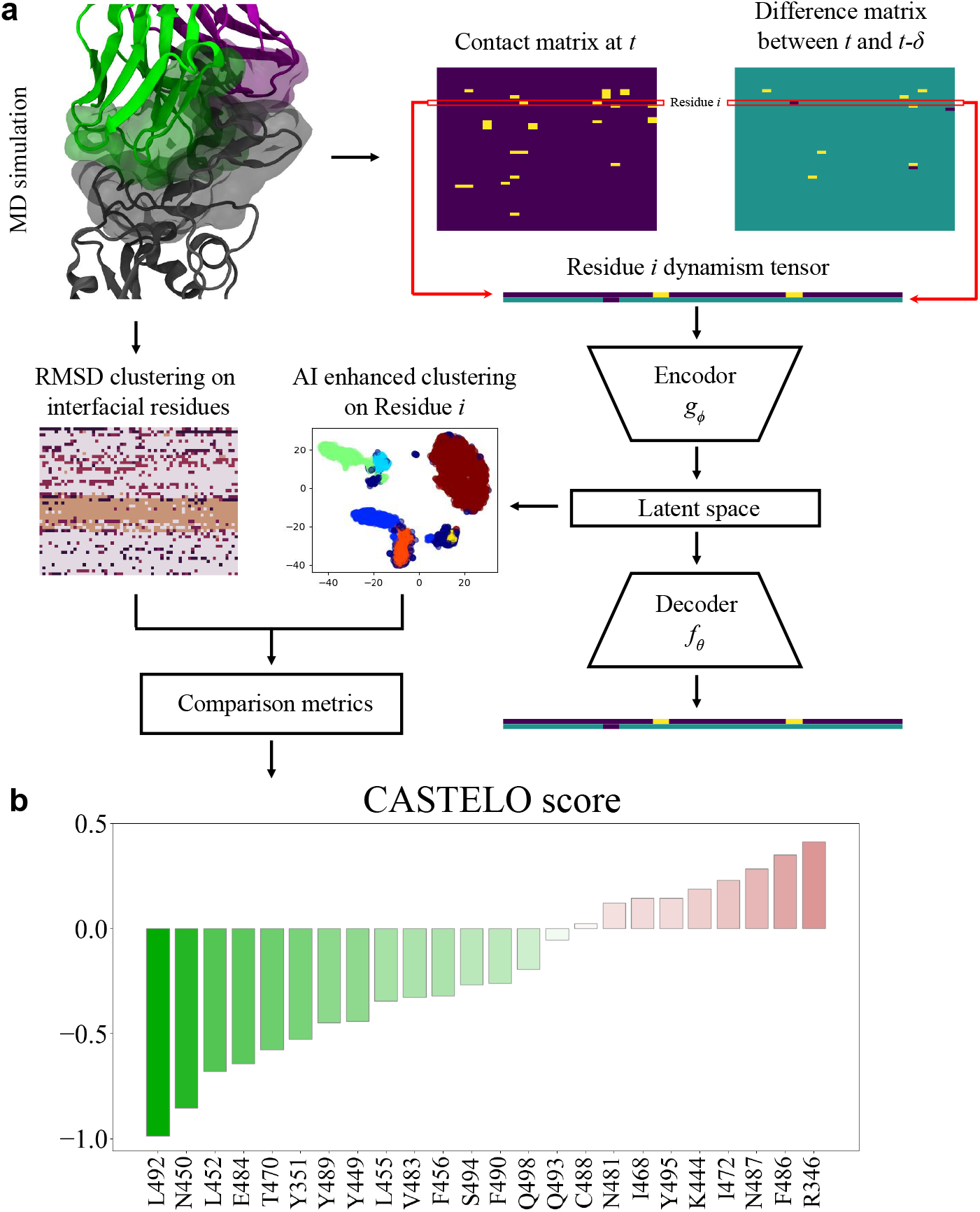
CASTELO workflow and CASTELO scores for the RBD/LY-CoV555 simulation. A: CASTELO workflow. We collect data from MD simulations. On one route, we calculate the clusters of the interfacial residues with RMSD clustering. On another route, we calculate the clusters of each residue *i* following these steps: (1) extracting contact matrix and difference matrix from the simulations; (2) assembling dynamism tensor for residue *i*; (3) compressing dimensions of the dynamism tensors with CVAE onto the latent space; (4) performing HDBSCAN clustering on the latent space for residue *i*. Finally, RMSD clustering results and AI-enhanced clustering results are compared with comparison metrics such as *dv* score. CASTELO score is calculated from the dv scores (using the equation in Methods). B: CASTELO scores are plotted for interfacial residues in RBD. The interfacial residues of RBD are defined as the residues in RBD that are within 5 Å of LY-CoV555 in its stable binding structure. Negative scores mean that the residues are beneficial. Positive scores mean that the residues are malicious.

We then calculated the contact matrix and dynamism tensor (see Methods) at every time frame *t* of the simulation (showing examples in Figure 4A). In order to find the clusters for individual residues, we used CVAE and HDBSCAN (see Methods section) on the calculated dynamism tensor at all time frames. Then, we utilize RMSD clustering results (using the interfacial residues) calculated above and CVAE clustering results calculated with dynamism tensors of the individual residues to evaluate the CASTELO scores for each of the interfacial residues (see Methods). The scores are plotted in Figure 4B, showing that L492, N450, L452 and E484 in RBD are among the four most beneficial residues to the complex of RBD and LY-CoV555 interactions. Intriguingly, from our MD simulation trajectory we found that L492 and L452 in RBD share close contacts with L55 of V*^H^* chain, which indicates an important hydrophobic core at the complex interface. Thus, similar to the L452R mutation, the possible L492R mutation can also reduce the LY-CoV555’s binding with RBD. N450 of RBD is physically close to this hydrophobic core, while sharing hydrogen bond through a water chain with S30 of V*^H^*. Thus, the mutation N450I (occurred with a single nucleotide variation in their codon codes) can disrupt this interfacial hydrophilic interaction and weaken the LY-CoV555’s binding with RBD. E484 of RBD, on the other hand, seems to stabilize the binding with a completely different mechanism by forming salt-bridges with R50 of V*^H^* and R96 of V*^L^*, as pointed out before. In addition, it is worth highlighting that CASTELO identified four other beneficial residues S494, Q493, F490 and V483 (Figure 4b) whose mutations were proven to be evasive for LY-CoV555 in experiment. ^22^

In contrast, R346, F486, N487 and I472 are among the four most malicious residues to the interactions between RBD and LY-CoV555. R346 of RBD was picked out, because it was highly flexible when being completely solvated in the electrolyte. Normally one should ignore a pick like this because it facilitates the solvation of the protein, even though this residue might be categorized as an interfacial residue by conventional standards. F486 of RBD forms an imperfect π – π stacking with slightly hydrophilic Y92 of V*^H^* in LY-CoV555, while being close to hydrophilic N487. As a result, N487 of RBD becomes unfavorable as well and was identified by CASTELO as a malicious residue. Finally, the hydrophobic I472 of RBD interestingly contacted with the highly hydrophilic E484-R50 and E484-R96 salt bridges. As a result, it was picked as an unstable residue by CASTELO. Therefore, any mutation among these four might increase the binding affinity between LY-CoV555 and RBD, which can be catastrophic for the viral survival in a host.

## Discussion and Conclusions

In this *in silico* study, we assumed that LY-CoV555 targets RBD-v at the same epitope, which leads to our conclusion that based on FEP calculations the double mutations L452R and E484Q can significantly reduce LY-CoV555’s binding affinity to RBD-v. Through MD simulation, we highlighted the molecular mechanism on how LY-CoV555 can be efficaciously bound with wide-type RBD but fails to bind RBD-v in variant B.1.617. Our alchemy FEP calculations show that binding free energy changes for L452R and E484K mutations are 3.04 and 22.22 kcal/mol respectively, indicating that these two mutations substantially weaken the binding between RBD-v and LY-CoV555 and thus are evasive. However, we emphasize that these two mutations are not resistant to LY-CoV016 (or Etesevimab that is another antibody in Lilly’s antibody cocktail), as shown in Figure S3 in the Supporting Information.

We employed CASTELO to further analyze what other residues can possibly weaken the RBD’s binding with LY-CoV555 after mutations. Interestingly, besides L452 and E484 whose mutations have occurred in B.1.617 already, CASTELO also identified L492 and N450 to be important for RBD’s binding with LY-CoV555. Noteworthily, L492 was found to be important for RBD’s binding with the antibody BD23,^25^ while N450 were previously identified as critical for the binding with the antibody CoV2-06^26^ and its mutations to other amino acids have proven to be resistant to various neutralizing antibodies.^27^ Here, possible L492R and N450I mutations in SARS-CoV-2 might be deleterious for LY-CoV555’s binding with RBD-v and thus can be evasive as well.

Besides LY-CoV555, these two mutations in the variant B.1.617 might reduce the binding affinity with other antibodies as well. For example, E484 coordinates N52/S55 (through hydrogen bonds) in the complex of RBD and the antibody S2M11, and consequently the mutation E484Q might weaken the complex, ^19^ which warrants further studies. Overall, our work shed light on the mechanism of evasive mutations of the variant B.1.617 and can facilitate the design of new antibody drugs specifically targeting the variant B.1.617.

## Methods

### MD simulations

We carried out all-atom MD simulations for the complex of RBD and the Fab of LY-CoV555 (PDB code: 7KMG) using the NAMD2.13 package^28^ running on the IBM Power Cluster. The complex was solvated in a cubic water box that measures about 110×110×110 Å^3^. K^+^ and Cl^−^ were added to neutralize the entire simulation system and set the ion concentration to be 0.15 M (Figs. 1a). The final simulation system comprises 132,308 atoms. The built system was first minimized for 10 ps and further equilibrated for 1000 ps in the NPT ensemble (*P* ~ 1 bar and *T* ~ 300 K), with atoms in the backbones harmonically restrained (spring constant *k*=1 kcal/mol/Å^2^). The production run was performed in the NPT ensemble without any restraint. The same approach was applied in the production run for RBD in a 0.15 M KCl electrolyte (Fig. S1 in Supporting Information), a free state required in the free energy perturbation (FEP) calculations (see below). The water box for the RBD-only simulation also measures about 110×110×110 Å^3^. Note that the same system size for the bound and the free states are required for free energy perturbation calculations for mutations with a net charge change.

We used the CHARMM36m force field^29^ for proteins, the TIP3P model^30,31^ for water, the standard force field^32^ for K^+^ and Cl^−^. The periodic boundary conditions (PBC) were applied in all three dimensions. Long-range Coulomb interactions were calculated using particle-mesh Ewald (PME) full electrostatics with the grid size of about 1 Å in each dimension. The pair-wise van der Waals (vdW) energies were computed using a smooth (10-12 Å) cutoff. The temperature *T* was kept at 300 K by applying the Langevin thermostat,^33^ while the pressure was maintained constant at 1 bar using the Nosé-Hoover method.^34^ With the SETTLE algorithm^35^ enabled to keep all bonds rigid, the simulation time-step was 2 fs for bonded and non-bonded (including vdW, angle, improper and dihedral) interactions, and the time-step for Coulomb interactions was 4 fs, with the multiple time-step algorithm. ^36^

### Free energy perturbation calculations

After equilibrating the structures in bound and free states, we performed free energy perturbation (FEP) calculations.^21^ In the perturbation method, many intermediate stages (denoted by λ) whose Hamiltonian *H*(λ)=λ*H_f_*+(1-λ)*H_i_* are inserted between the initial and final states to yield a high accuracy. With the softcore potential enabled, *λ* in each FEP calculation for the bound or free state varies from 0 to 1.0 in 20 perturbation windows (lasting 300 ps in each window), yielding gradual (and progressive) annihilation and exnihilation processes for L452 and R452 (or for E484 and Q484), respectively. More detailed procedures can be found in our previous work.^18?^ In FEP runs for the E484Q mutation, the net charge of the MD system changed from −1 to 0 e (where e is the elementary charge). It is important to have similar sizes of the simulation systems for the free and the bound states,^37,38^ so that the energy shifts from the Ewald summation (due to the net charge in the final simulation system) approximately cancel out when calculating ΔΔ*G*.

### CASTELO Pipeline

CASTELO,^23^ or “clustered atom subtypes aided lead optimization”, is a combined molecular dynamics (MD) and machine learning (ML) method that identifies improvable atoms or atom groups in a drug lead molecule (such as small molecule drugs^23^ and peptides^24^). We followed the previous study, ^24^ where CASTELO was applied to a protein-peptide binding simulations and applied the same workflow on the RBD-antibody binding simulations. In short, we calculated the contact matrix between RBD and antibody at each individual timeframe (time *t*) of the simulations. The elements of the calculated contact matrix consisted of the closest-heavy distances between *M* residues of antibody and *N* residues of the RBD. Note that closest-heavy distance between residue *i* and residue *j* is calculated as the minimal distance between all interfacial pairs of heavy atoms (non-hydrogen atoms) in these two residues. In addition to the contact matrix, we enriched the extracted information at time *t* with the difference contact matrix between *t* and *t* – *δ*. The resulting 2 x *M* x *N* matrix is coined “dynamism tensor”, which contains both static contact information and dynamic information of the protein binding complex at time *t*. Subsequently, we used Convolutional Variational Autoencoders (CVAE) to represent each dynamism tensor in a *d*-dimensional latent space (with a vector of *d* elements). For each residue, we used HDBSCAN^39^ as our clustering method to identify the clusters at the latent space (Figure 4A). As a comparison, we used conventional RMSD clustering method (quality threshold method in VMD^40^) to identify the clusters of the binding interface (defined as the residues within 5 Å of its binding partner). Eventually, we compared the individual residue-wise clusters and the binding interface clusters with comparison metrics (summarized as the CASTELO score, plotted in Figure 4B) to rank the residues from most beneficial to the protein-protein interactions to the most malicious to the interactions. To calculate the CASTELO scores, we first utilize the dv scores presented by Gionis et al.^41^ that compares two clustering results (see Bell et al.^24^). We then normalize the dv scores using the equation: (*dv_i_* – ∑*_i_dv*/*N*) /max*_i_*(*dv_i_* – ∑*_i_dv*/*N*), which is denoted as the CASTELO score. All parameters were adopted from a previous study, ^24^ except that the cutoff distance of the contact matrix was changed to 4 Å and convolutional filter size was changed to 1 x 3 to accommodate the small cutoff distance.

## ASSOCIATED CONTENT

### Supporting Information

An illustration of buried salt-bridges inside the complex; MD simulation of the RBD-only system; A movie showing the stable interfacial salt-bridge binding (E484-R50-R96.mpg); A movie showing the flexible coil (containing E484) in the RBD-only simulation.

### Notes

T. H., L.Z. and B. L. declare no competing financial interest.

## ACKNOWLEDGEMENT

T.H., L.Z. and B.L. gratefully acknowledge the computing resource from the IBM Cognitive Computing Program.

## Notes

### Competing Interest Statement

The authors have declared no competing interest.

